# Evidence against altered excitatory/inhibitory balance in the posteromedial cortex of young adult APOE E4 carriers: a resting state ^1^H-MRS study

**DOI:** 10.1101/2021.05.12.443879

**Authors:** AG Costigan, K Umla-Runge, CJ Evans, R Raybould, KS Graham, AD Lawrence

## Abstract

A strategy to gain insight into early changes that may predispose people to Alzheimer’s disease is to study the brains of younger cognitively healthy people that are at increased genetic risk of AD. The Apolipoprotein (APOE) E4 allele is the strongest genetic risk factor for AD, and several neuroimaging studies comparing APOE E4 carriers with non-carriers at age ~20-30 have detected hyperactivity (or reduced deactivation) in posteromedial cortex (PMC), a key hub of the default network (DN) which has a high susceptibility to early amyloid deposition in AD. Transgenic mouse models suggest such early network activity alterations may result from altered excitatory/inhibitory (E/I) balance, but this is yet to be examined in humans. Here we test the hypothesis that PMC fMRI hyperactivity could be underpinned by altered levels of excitatory (glutamate) and/or inhibitory (GABA) neurotransmitters in this brain region. Forty-seven participants (20 APOE E4 carriers and 27 non-carriers) aged 18-25 underwent resting-state proton magnetic resonance spectroscopy (^1^H-MRS), a non-invasive neuroimaging technique to measure glutamate and GABA *in vivo*. Metabolites were measured in a PMC voxel of interest and in a comparison voxel in the occipital cortex (OCC). There was no difference in either glutamate or GABA between the E4 carriers and non-carriers in either MRS voxel, nor in the ratio of glutamate to GABA, a measure of E/I balance. Default Bayesian t-tests revealed evidence in support of this null finding. Results suggest that PMC hyperactivity in APOE E4 carriers is unlikely to be associated with, or indeed may precede, alterations in local resting-state PMC neurotransmitters, thus informing the spatio-temporal order and the cause/effect dynamic of neuroimaging differences in APOE E4 carriers.

**Highlights:**

- Hyperactivity in posteromedial (PM) network in people at AD genetic risk (APOE E4)
- Such PM network hyperactivity may initiate pathogenic cascade that triggers AD
- APOE mouse models suggest hyperactivity driven by excitatory/inhibitory imbalance
- Using ^1^H-MRS at 3T we studied PMC E/I balance in young adult APOE E4 carriers
- Found evidence against altered E/I balance in young adult APOE E4 carriers

## 1 Introduction

The Apolipoprotein (APOE) E4 allele is the strongest genetic risk factor for late onset Alzheimer’s disease (LOAD), where possession of one E4 allele increases risk of AD by 3-4 times, and two alleles by 12-14 times compared to the AD-neutral E3/E3 genotype (Belloy et al., 2019; Farrer et al., 1997). There is also a gene-dose effect on the age of onset of possible AD, where possession of 0, 1 and 2 E4 alleles reduces age of onset from 84 to 76 to 68 years, respectively (Corder et al., 1993; van der Lee et al., 2018). The mechanisms by which APOE increases AD risk are not clear. There is a growing consensus that AD-related brain changes and pathology occur decades before the onset of symptoms (Jack et al., 2018; Jagust, 2018; Masters et al., 2015; Sperling et al., 2014). By comparing APOE E4 carriers and non-carriers decades before the typical AD onset age, and presumably free from AD pathology, we can gain insight into differences which may predispose APOE E4 carriers to developing AD.

The posteromedial cortex (PMC) is a key region of interest for such studies in young people at AD genetic risk. The PMC (including retrosplenial cortex (RSC), posterior cingulate cortex (PCC) and precuneus (PCu)) (Parvizi et al., 2006) constitutes a major cortical hub that is densely connected with the medial temporal lobe to form a corticohippocampal brain network, referred to as the posterior medial (PM) network, a subsystem of the default network (DN) (Raichle, 2015; Ranganath & Ritchey, 2012). The PM network is critical to episodic memory and related cognitive processes relevant to AD (Ranganath & Ritchey, 2012; Ritchey & Cooper, 2020). The PMC is particularly susceptible to early amyloid (Aβ) plaque deposition, one of the hallmark pathologic features of AD (Buckner et al., 2005; Maass et al., 2019; Palmqvist et al., 2017), with APOE E4 allele carriers having both a younger age of onset and faster rates of PMC amyloid deposition relative to non-carriers (Burnham et al., 2020; Mishra et al., 2018). The cause of early Aβ aggregation within PMC is currently unknown, but may reflect a lifespan regional vulnerability (Buckner et al., 2005).

Aberrant PMC network activity is consistently seen in individuals at high risk for AD and during the early stages of the disease. Functional magnetic resonance imaging (fMRI) studies reveal reduced PMC deactivation in conjunction with hippocampal hyperactivation during episode encoding tasks (see Palop & Mucke, 2016; Zott et al., 2018 for review). Inadequate deactivation of the PMC is associated with amyloid deposition and poorer episodic memory performance, and has been related to conversion from mild cognitive impairment (MCI) to AD (Palop & Mucke, 2016; Zott et al., 2018). In later stages of AD, the reduced DN deactivation may persist, whereas the hippocampus is hypoactive during memory encoding (Palop & Mucke, 2016; Zott et al., 2018).

Such PMC functional alterations, which mirror the functional signature of early AD, have been found to begin decades before the clinical onset of AD in participants at increased AD genetic risk. For example, fMRI studies in young adult APOE E4 carriers have detected lack of PMC/DN deactivation and hippocampal hyperactivity in carriers compared to non-carriers during episodic encoding and spatial memory tasks (Filippini et al., 2009; Kunz et al., 2015; Persson et al., 2008; Shine et al., 2015). Furthermore, fMRI studies in participants with familial AD (FAD) mutations (Presenilin1, PSEN1) have detected reduced DN deactivation decades before the typical age-of-onset of AD (Mondadori et al., 2006; Reiman et al., 2012). As the functional alterations in the PM network in people at risk of developing AD widely overlap with regions that ultimately develop AD pathology and hypometabolism/atrophy, this hyperactivity may be a harbinger and even a cause of AD rather than a compensatory phenomenon (Buckner et al., 2005; Jagust & Mormino, 2011; Palop & Mucke, 2016).

Indeed, AD mouse models provide evidence that hyperactivity is causally linked to/drives amyloid deposition. In APP transgenic mice expressing a mutated form of Aβ precursor protein, lactate (as a measure of neuronal activity) was closely associated with interstitial fluid (ISF) Aβ levels, and in turn, ISF Aβ predicted region-specific Aβ deposition, particularly in PMC (Bero et al., 2011). In addition, chronic optogenetic activation in young FAD mice promotes amyloid deposition (Yamamoto et al., 2015). A recent study of MCI patients provided human evidence that hyperactivity is associated with subsequent amyloid deposition (Leal et al., 2017). Collectively, these studies support the hypothesis that Aβ accumulates in a concentration-dependent manner throughout life, with selective neuronal and network hyperactivity leading to excessive Aβ production, ultimately causing aggregation and seed formation that then spreads trans-synaptically, following the path of intrinsic brain networks (Bero et al., 2011; Jagust & Mormino, 2011; Seeley, 2017; Sepulcre et al., 2018).

It is important, therefore, to better understand the PMC hyperactivity detected in young APOE E4 carriers, as this could provide important insight into the earliest changes and timeline of changes that may predispose to AD. One such mechanism that may underpin PMC hyperactivity is alterations in the levels of the inhibitory neurotransmitter, γ-aminobutyric acid (GABA), and excitatory neurotransmitter, glutamate. The balance between these neurotransmitters, termed the excitatory/inhibitory (E/I) balance, mediates normal neural network activity (Busche & Konnerth, 2016; Palop & Mucke, 2016). A growing consensus views AD as a circuit-based disorder, where an alteration of the physiological E/I balance underlies both the functional impairment of local neuronal circuits as well as that of large-scale networks in the amyloid-depositing brain (Busche & Konnerth, 2016; Harris et al., 2020; Palop & Mucke, 2016). Mouse models of AD suggest that a shift in E/I balance towards excitation initially causes hyperactivity in cortical and hippocampal neurons, prior to the appearance of amyloid plaques (with hypoexcitability in late disease stages, linked to subsequent tau deposition) (Bi et al., 2020; Busche & Konnerth, 2016; Harris et al., 2020; Jiménez-Balado & Eich, 2021; Najm et al., 2019; Palop & Mucke, 2016).

The APOE E4 genotype may be a distinct risk factor for hyperexcitability (Toniolo et al., 2020), linked both to an increase in excitatory tone and a decrease in GABAergic inhibition. For example, a study combining fMRI with electrophysiology identified that entorhinal cortex hyperactivity in APOE E4 mice, in the absence of overt AD pathology, was associated with decreased response to inhibitory GABAergic inputs on pyramidal neurons, rather than increased excitability due to differences in NMDA glutamate receptors (Nuriel et al., 2017). Alternatively, impairment in glutamate production in APOE E4 transgenic mice has been suggested, as E4 compared to E3 mice had a lower concentration of glutamate and glutaminase, the enzyme that converts glutamine to glutamate, and a higher concentration of glutamine (Dumanis et al., 2014). Thus, the levels of neurotransmitters in young APOE E4 carriers could provide important insight into why the PMC demonstrates hyperactivity, and why this is a key region affected early by amyloid pathology (Burnham et al., 2020).

In human participants, the concentration of neurotransmitters can be measured non-invasively *in vivo* using proton magnetic resonance spectroscopy (^1^H-MRS) (Stagg & Rothman, 2014). Glutamate and GABA+ (GABA plus co-edited macromolecules; see Section 2.5) quantified via ^1^H-MRS in the PMC have been associated with the magnitude of the local BOLD response, showing that regional excitation-inhibition balance predicts default network deactivation. In these PMC studies, higher concentrations of inhibitory GABA+ were associated with greater task-related deactivation, higher levels of excitatory Glx with less deactivation, and a higher E/I (i.e. Glx/GABA+) ratio with less deactivation (Gu et al., 2019; Hu et al., 2013). An unexplored avenue is whether the PMC hyperactivity/reduced deactivation in young APOE E4 carriers could reflect altered GABA+ and or Glx levels and a shift in E/I balance toward excitation, as suggested by animal models.

^1^H-MRS studies of E/I neurotransmission in older individuals with APOE E4, MCI and AD have produced inconsistent results, and are confounded by the presence of AD pathology and other age-related brain changes. Several studies show that AD and MCI individuals have lower PMC GABA and Glx than age-matched healthy controls (Bai et al., 2014; Londono et al., 2013; Oeltzschner et al., 2019; Riese et al., 2015). It is less clear whether older APOE E4 carriers have altered PMC GABA+ and Glx levels prior to amyloid pathology: E4 carriers appeared to have lower PMC GABA+ and Glx compared to non-carriers at approx. age 70, however this was based on a small sample size of nine E4 carriers and was not statistically significant (Riese et al., 2014). PMC Glx was compared in a larger sample of E4 carriers and non-carriers at ages 20-70 (split into young and old groups). No difference in PMC Glx was found between APOE groups when collapsed across age groups (Suri et al., 2017). A recent published abstract found that higher levels of GABA and Glu were associated with higher amyloid burden in a group of 30 older individuals, which was particularly evident in APOE E4 carriers, but this study did not examine GABA/Glu (E/I) ratio (Schreiner et al., 2016). Two previous studies have investigated other ^1^H-MRS metabolites in the PMC in young APOE E4 carriers (Calderón-Garcidueñas et al., 2015; Suri et al., 2017; see Supplementary Material, Section 6 for replication), however this study is the first to measure both Glx and GABA+, and the E/I balance (Glx:GABA+ ratio) in the PMC using ^1^H-MRS.

Based on consistent findings of reduced PMC deactivation in young APOE E4 carriers, and evidence suggesting this is linked to an E/I imbalance, we hypothesized that young APOE E4 carriers would show a shift in E/I balance toward excitation due to either/both lower PMC GABA+ and higher Glx than non-carriers. This will be reflected by an increased PMC E/I (Glx:GABA+) ratio in E4 carriers compared to non-carriers. To assess regional specificity of any difference, as recommended in Duncan et al (2014), we used a comparison voxel, placed outside the PMC in the occipital cortex (OCC). The OCC is a region not reported to show altered BOLD in fMRI studies of young APOE E4 carriers relative to non-carriers (Filippini et al., 2009; Kunz et al., 2015; Shine et al., 2015).

To assess the extent to which our data set provides evidence for or against our research hypothesis, we use Bayes factors (Dienes & Mclatchie, 2018; Morey et al., 2016). Bayes factors (BFs) “allow accepting and rejecting the null hypothesis to be put on an equal footing” (Dienes, 2014), with evidence either for or against an E/I imbalance toward excitation in young adults at increased genetic risk for AD being of considerable evidential value for understanding the temporal sequence of early network dysfunction underlying APOE E4 related AD risk (Harris et al., 2020; Jagust & Mormino, 2011; Palop & Mucke, 2016).

## 2 Materials and Methods

### 2.1 Participants

Participants were recruited from two cohorts of undergraduate Psychology students who provided a saliva sample for APOE-genotyping. The first cohort (total n=125) were the same as that in Shine et al. (2015), and Hodgetts et al. (2019), and the second cohort comprised a further new 229 students. Nineteen participants (10 APOE-E4 carriers and 9 non-carriers) from the first cohort and 28 (10 APOE-E4 carriers and 18 non-carriers) from the second cohort took part in this study. The total sample size, therefore, was 20 APOE-E4 carriers and 27 non-carriers (see Table 1). There was one male in each group, reflecting the predominantly female undergraduate Psychology cohort.

**Table 1:** Demographic details of the APOE E4 carrier and non-carrier groups, and number of participants scanned in each group.

|  | APOE E4 carriers | Non-carriers |
| --- | --- | --- |
| Total n | 20 | 27 |
| Genotype split | 1 E4/E4, 18 E3/E4, 1 E2/E4 | 24 E3/E3, 3 E2/E3 |
| N females/males | 19 / 1 | 26 / 1 |
| Age (mean ± SD) | 21.2 ± 1.9 | 19.9 ± 1.6 |

APOE-E4 carriers and non-carriers were matched for age, family history of dementia, and family history of psychiatric illness. Participants were excluded if they had a self-reported history of depression or psychiatric illness or were taking any psychoactive medication. All participants were right-handed, with normal or corrected-to-normal vision.

A double-blind strategy was adopted for this study, whereby both participants and researchers collecting and analysing data were blind to the participants’ APOE-genotypes, in order to prevent any bias during analyses. The study received ethical approval from the Cardiff University School of Psychology Research Ethics Committee, and all participants provided written informed consent.

The sample size was estimated via a priori power calculation, using G*Power version 3.1.9.7 (Faul et al., 2009). Based on the fMRI study of Shine et al. (2015) in a similarly aged population, which detected a very large effect size (Cohen’s d > 1) for the difference in PMC deactivation between APOE E4 carriers and non-carriers, we planned our experiment to detect a conventionally large effect (Cohen’s d = 0.8) while safeguarding against effect size inflation in published studies (Perugini et al., 2014, 2018). We initially aimed for a sample size of n = 26 per group, which would have 80% power to detect such a large effect at p < 0.05 using a 2-tailed t-test. The final obtained sample of 20 APOE E4 carriers and 27 non-carriers was similar to previous ^1^H-MRS studies in young APOE E4 carriers measuring other metabolites (tNAA, mI, Cho, tCr), in which the sample sizes were 8 APOE E4 carriers vs 22 non-carriers (Suri et al., 2017), and 22 *APOE-E4* carriers vs 28 non-carriers (Calderon-Garciduenas et al., 2015), as well as a study of GABA and glutamate in older individuals without cognitive impairment (Quevenco et al., 2019).

### 2.2 APOE genotyping

Procedures for DNA extraction from saliva and *APOE* genotyping were the same for the two cohorts. DNA was obtained from saliva using Oragene OG-500 saliva kits (DNA Genotek, Inc., Ontario, Canada). DNA extraction and *APOE*-genotyping were performed in the Centre for Neuropsychiatric Genetics and Genomics at Cardiff University. Since *APOE* isoforms differ due to a single nucleotide polymorphism (SNP) at two sites in the gene, a single SNP genotyping assay was performed for each site to determine *APOE* genotype. The SNP rs429358 was determined by KASP genotyping and rs7412 by Taqman genotyping. These were detected on Tecan infinite F200 pro and StepOnePlus^™^ Real-Time PCR System platforms, respectively. Haplotypes corresponding to *APOE* E2, E3 and E4 were then deduced.

Genotyping was successful in 100/125 and 224/229 participants from the two cohorts respectively. The distribution of genotypes of those successfully genotyped in the first cohort was E2/E2 (1/100, 1%), E2/E3 (10/100, 10%), E2/E4 (1/100, 1%), E3/E3 (69/100, 69%), E3/E4 (19/100, 19%), and E4/E4 (0/100, 0%). The genotype-distribution in the second cohort was E2/E2 (0/224, 0%), E2/E3 (38/224, 17%), E2/E4 (7/224, 3%), E3/E3 (125/224, 56%), E3/E4 (52/224, 23%), and E4/E4 (2/224, 1%).

### 2.3 MRI scan acquisition

All scans were performed at the Cardiff University Brain Research Imaging Centre (CUBRIC) on a 3T General Electric (GE) HDx scanner fitted with an 8-channel phased array head coil. A high resolution anatomical MRI scan was obtained for each participant using a 3D T1-weighted (T1w), fast spoiled gradient echo (FSPGR) sequence (TE/TR = 3.0/7.9ms; TI=450ms; flip angle 20°; data matrix 256×192×176; field of view 256×192×176mm^3^; acquisition time approx. 7 minutes). The FSPGR was used to aid ^1^H-MRS voxel placement during scanning (see Section 2.5).

The phase of menstrual cycle has been suggested to have an impact on ^1^H-MRS metabolite concentrations (Batra et al., 2008; De Bondt et al., 2015; Epperson et al., 2002), therefore scans of female participants were scheduled during their luteal phase (days 15-28 of the cycle, where day 1 was defined as the first day of menstruation). No restrictions were placed on scan scheduling if female participants were taking the contraceptive pill, as GABA+ concentration does not significantly differ between pill-on and pill-free days (De Bondt et al., 2015).

### 2.4 ^1^H-MRS acquisition

Single voxel proton spectra were acquired from the PMC (the voxel of interest, measuring 2×2×3cm^3^), and the occipital cortex (OCC, the comparison voxel, measuring 3×3×3cm^3^) in the resting-state. Examples of voxel placement are shown in Figure 1.

**Figure 1:**
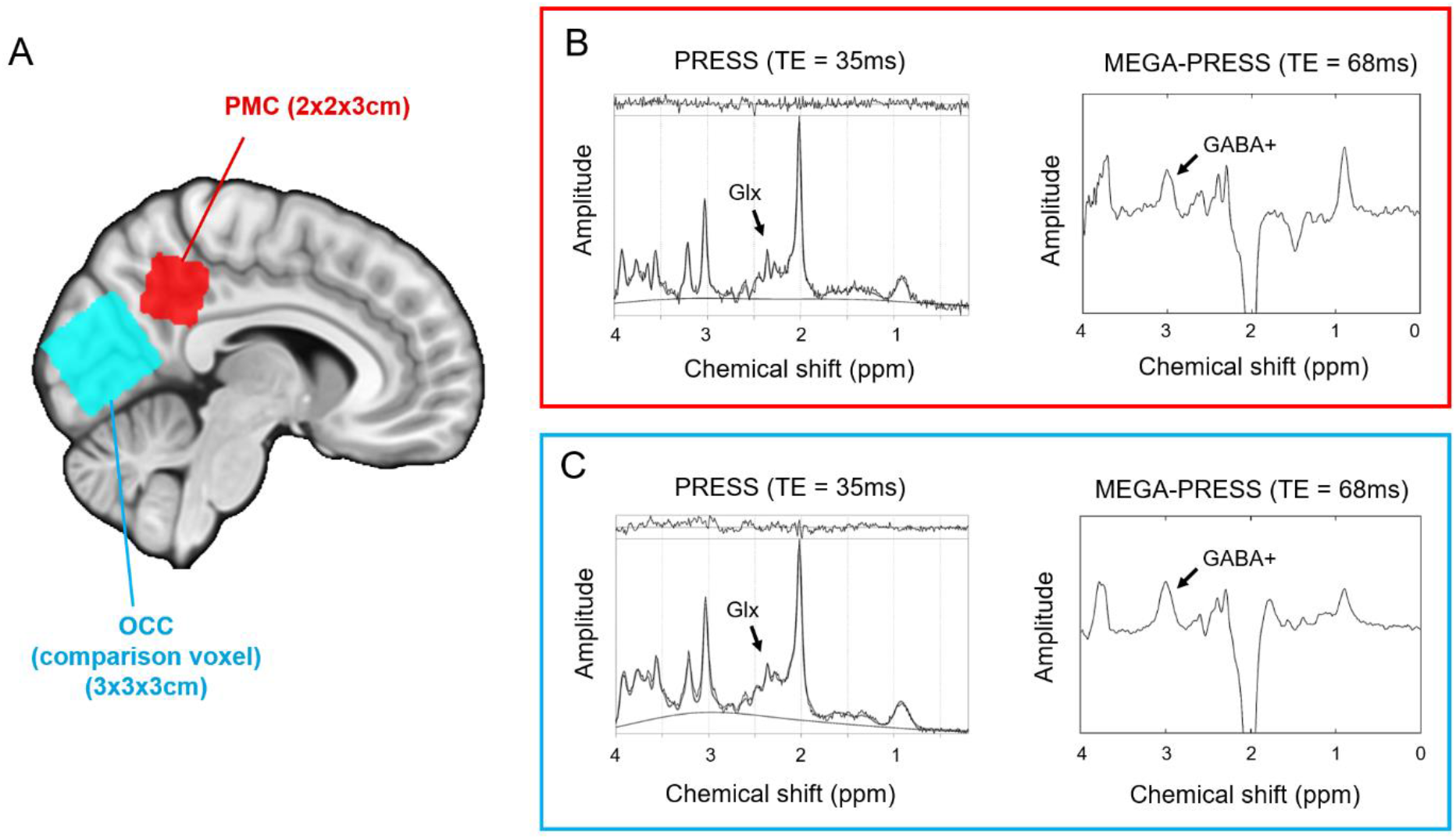
1H-MRS voxel placement and 1H-MRS spectra. (A) ^1^H-MRS voxel placement for one representative participant. Voxels have been transformed into MNI standard space and overlaid on the MNI152 2 mm standard brain template (Grabner et al., 2006). Example of a PRESS and MEGA-PRESS spectrum from (B) the PMC voxel and (C) OCC voxel in one participant.

Landmarks used for voxel placement were consistent with Costigan et al. (2019), which was an approach developed from a pilot study assessing test-retest reliability of voxel placement and metabolite concentrations (see Supplementary Material, Section 5). Briefly, an odd number of AC-PC aligned slices were acquired from the bottom to the top of the corpus callosum (typically 5 or 7 slices). The PMC voxel was placed on the plane of the middle slice, and adjusted to lie posterior to the ventricles, ensuring it was not covering an area of CSF to prevent artefacts in the ^1^H-MRS spectrum. The OCC voxel was placed above the line of the tentorium cerebelli and adjusted so it did not contain any scalp tissue, which would have resulted in lipid contamination in the spectra.

In each voxel, one point-resolved spectroscopy (PRESS) scan was obtained to measure Glx (TE/TR = 35/1500ms; number of averages = 128; scan time 4mins) (Bottomley, 1984). One Mescher-Garwood PRESS (MEGA-PRESS) scan (Mescher et al., 1998; Rothman et al., 1993) was obtained to measure GABA + coedited macromolecules, “GABA+”, (TE/TR = 68/1800ms; OCC 166 edit on/off pairs, scan time 10mins; PMC 256 on/off pairs, scan time 15mins). Shimming was performed before all ^1^H-MRS scans to ensure water-linewidth of 10Hz or lower, in order to obtain sharp peaks in the resulting ^1^H-MRS spectrum.

Different MRS sequences were used to quantify Glx and GABA+ so that the most optimal method was used for each metabolite. The MEGA-PRESS spectral editing acquisition was necessary because GABA+ is challenging to quantify accurately using standard PRESS methods, due to its very low concentration, and its peak resonances in the MRS spectrum have a low amplitude and overlap with other metabolites (mainly by creatine at 3.0ppm) (Harris et al., 2017; Mullins et al., 2014). MEGA-PRESS acquisitions include additional editing pulses placed symmetrically about the water resonance (4.7ppm) resulting in editing pulses at 1.9ppm (edit on) and at 7.5ppm (edit off) in order to subtract the creatine peak, enabling accurate GABA+ detection and quantification.

A separate PRESS sequence was used to quantify Glx, rather than using the MEGA-PRESS edit off scan as used in some studies, because short TE PRESS scans (e.g. TE 35ms, rather than TE 68ms as in the MEGA-PRESS edit off scan) have been found to be more accurate for our voxel of interest. For example, a test-retest reliability study found the 35ms PRESS scan produced a lower coefficient of variation and lower Cramer-Rao Lower Bounds (CRLBs) for PMC Glx over three intra-scan repeats than PRESS scans with a longer TE (Hancu, 2009).

The difference in voxel sizes was a trade-off between voxels being large enough to have a good signal-to-noise ratio (SNR) for accurate metabolite quantification, yet not too large, to maintain spatial specificity. The 3×3×3cm OCC MEGA-PRESS protocol has reliably produced good quality spectra to quantify GABA+ (e.g. Muthukumaraswamy et al., 2009; Muthukumaraswamy, Evans, Edden, Wise, & Singh, 2012) and has been recommended for MEGA-PRESS scanning (Mullins et al., 2014). A PMC voxel of this size, however, would have reduced the spatial specificity of our region of interest, and also overlapped with the OCC voxel. As the spatial specificity of the PMC voxel was of key importance here, its size was reduced to 2×2×3cm, which is more consistent with the PMC voxel size employed in previous studies of APOE carriers, and MCI or AD patients (Kantarci et al., 2000, 2002; Suri et al., 2017; Voevodskaya et al., 2016). To counteract the reduction in SNR from this volume reduction, the length of the PMC MEGA-PRESS scan was increased to make the SNR similar across the two MRS voxels.

### 2.5 Data analysis

PRESS data were analysed using TARQUIN (Totally Automatic Robust Quantification In NMR) version 4.3.3 (Wilson et al., 2011). The largely overlapping resonances of glutamate and glutamine make it difficult to accurately separate their spectra, therefore these measures were combined to create a composite glutamine + glutamate measure, or “Glx” (Rae, 2014; Stagg & Rothman, 2014). Glx data were excluded if the CRLB was above 20%, consistent with the data quality criteria recommended in the ^1^H-MRS literature (Cavassila et al., 2001; Lin et al., 2021; Near et al., 2020).

MEGA-PRESS data were analysed using GANNET (GABA-MRS Analysis Tool) version 2.0 (Edden et al., 2014). The GABA concentration measured in the MEGA-PRESS scans represents GABA plus co-edited macromolecules, and is referred to as “GABA+” (Mullins et al., 2014). Data quality was assessed by two independent raters (authors AGC and CJE) using a 3-point rating scale (very good, satisfactory, reject; as in Lipp et al., 2015). Inter-rater reliability was assessed via the coefficient of variation (CV), which found good correspondence between raters for both PMC (CV=8.46%) and OCC (CV=8.27%).

Metabolite concentrations were corrected for voxel composition: using each participant’s high resolution T1w anatomical MRI scan, FSL’s FAST tool segmented the areas of the PMC and OCC ^1^H-MRS voxels into cerebrospinal fluid (CSF), grey matter (GM) and white matter (WM) (Zhang et al., 2001). Metabolites were quantified using the tissue H_2_O signal as an internal concentration reference and are expressed as a concentration in millimoles (mM) per unit tissue volume (Near et al., 2020). The water signal was preferable here, as opposed to using tCr as the reference metabolite, to prevent ambiguity for whether there were alterations in the numerator (i.e. Glx or GABA+) or denominator (tCr) between APOE carriers and non-carriers (Wilson et al., 2019). Furthermore the water signal has a much higher signal that that of the tCr peak, therefore is more reliable for quantification (Near et al., 2020). The metabolite signals were corrected for the proportion of CSF in the voxel (as the concentration of metabolites in CSF is negligible) and the water reference signal was corrected to account for the differing water content of CSF, GM and WM (Near et al., 2020; Wilson et al., 2019).

### 2.6 Statistics

The metabolites Glx, GABA+ and the ratio of Glx:GABA+ (as a measure of E/I balance (Steel et al., 2020)) were compared between APOE E4 carriers and non-carriers in each voxel using two-tailed independent sample t-tests in SPSS version 26. Cohen’s d was calculated as a measure of effect size. To support the frequentist statistics, Bayesian t-tests were implemented in JASP version 0.13.1. The Bayes factor (BF) assesses the strength of the evidence that the data provide for the null hypothesis (H0, i.e. no difference between APOE groups) versus the alternative hypothesis (H1, i.e. difference between groups), expressed as BF_01_, or for the alternative versus the null hypothesis, denoted BF10. Where the frequentist t-test was non-significant, BF_01_ is reported to support this null finding. Bayes factors grade the strength of evidence on a continuous scale (with a BF_01_ of 1 indicating the finding is equally likely under H0 and H1), although a BF_01_ value over 3 is frequently interpreted as substantive evidence for the null hypothesis (Hu et al., 2018; Keysers et al., 2020). Graphs were created using GraphPad Prism version 5.01 for Windows, GraphPad Software, San Diego California USA, www.graphpad.com.

## 3 Results

### 3.1 Participants and data quality exclusions

The genotype split of the 20 APOE E4 carriers and 27 non-carriers is shown in Table 1. There was no significant difference in age between groups, (t(45)=1.14, p=0.26, Cohen’s d=0.34)), nor in the proportion of males in each group (Fisher’s exact test p=1.00 (2 tailed)).

Data quality assessments resulted in the exclusion of three PMC Glx (one E4 carrier) and two OCC Glx scans (two E4 carriers), and 14 PMC GABA+ (six E4 carriers) and one OCC GABA+ (E4 carrier) scans. One PMC GABA+ scan was excluded as an outlier (GABA+ concentration was 4 standard deviations from the mean). Final sample sizes for each metabolite in the two MRS voxels are shown in Table 2.

**Table 2:** Metabolite results, sample sizes and summary of statistical analysis. Metabolite values are shown as the mean concentration (mM) and standard error of the mean. Sample sizes indicate the number of good quality data sets remaining after data quality assessment and outlier exclusion. Statistical tests are a 2-tailed t-test and a 2-tailed Bayesian t-test to assess the strength of the evidence for the null hypothesis (i.e. no difference between groups, denoted BF_01_).

| MRS voxel | PMC |  |  |  | OCC |  |  |  |
| --- | --- | --- | --- | --- | --- | --- | --- | --- |
|  | E4 carriers | Non-carriers | t-test p value | BF<sub>01</sub><br>(evidence for null) | E4 carriers | Non-carriers | t-test p value | BF<sub>01</sub><br>(evidence for null) |
| **Glx** | 20.86 | 21.21 | 0.75 | 3.22 | 21.01 | 20.59 | 0.75 | 3.24 |
| St err | 0.82 | 0.74 |  |  | 0.92 | 0.87 |  |  |
| n | 19 | 26 |  |  | 19 | 27 |  |  |
| **GABA+** | 2.07 | 2.03 | 0.83 | 2.90 | 1.91 | 1.96 | 0.38 | 2.47 |
| St err | 0.13 | 0.45 |  |  | 0.03 | 0.05 |  |  |
| n | 14 | 18 |  |  | 19 | 27 |  |  |
| **Glx : GABA+** | 10.67 | 10.61 | 0.96 | 2.93 | 11.08 | 10.62 | 0.54 | 2.89 |
| St err | 0.86 | 0.59 |  |  | 0.52 | 0.50 |  |  |
| n | 14 | 17 |  |  | 19 | 27 |  |  |

### 3.2 Regional specificity of PMC and OCC metabolites

There were no significant correlations between PMC metabolites and OCC metabolites, suggesting regional specificity of metabolite concentrations: PMC vs OCC Glx, r(45)=0.22, p=0.15, BF_01_=1.97; PMC vs OCC GABA+, r(33)=0.13, p=0.49, BF_01_=3.66; PMC vs OCC Glx:GABA+, r(33)=0.22, p=0.22., BF_01_=2.22.

### 3.3 Glx

There was no statistically significant difference in PMC Glx between APOE E4 carriers and non-carriers: t(43)=0.32, p=0.75; Cohen’s d= −0.10; BF_01_ = 3.22. There was also no statistically significant difference in Glx between APOE groups in the OCC comparison voxel: t(44)=0.32, p=0.75; Cohen’s d=0.10, BF_01_ = 3.24 (see Figure 2A).

**Figure 2:**
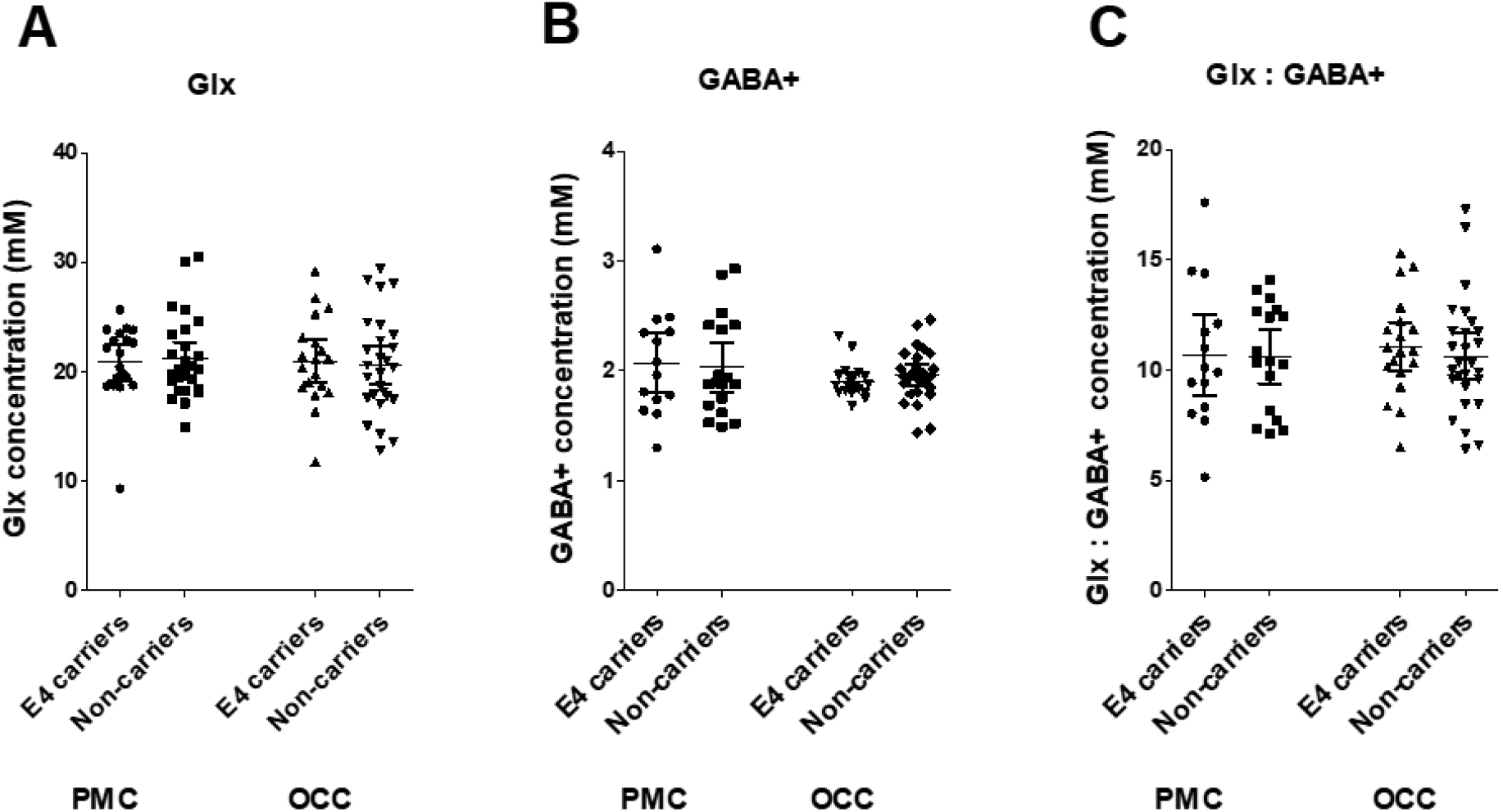
Comparison of metabolite concentrtaions in APOE-E4 carriers and non-carriers in the voxel of interest, the PMC, and the comparison voxel in the OCC. (A) Glutamate (Glx); (B) GABA+; (C) the ratio of glutamate to GABA+ as a measure of excitatory/inhibitroy (E/I) balance. Each dot represents one participant, the horizontal lines represent the mean and the error bars are 95% confidence intervals.

### 3.4 GABA+

There was no statistically significant difference in PMC GABA+ between E4 carriers and non-carriers: t(30)=0.22, p=0.83; Cohen’s d=0.08, BF_01_ = 2.90. There was also the case for GABA+ in the OCC comparison voxel: t(44)=0.88, p=0.38, Cohen’s d= −0.26, BF_01_ = 2.47 (See Figure 2B).

### 3.5 Excitatory/Inhibitory (E/I) balance (Glx/GABA+)

There was no statistically significant difference in the ratio of PMC Glx:GABA+ between E4 carriers and non-carriers: t(29)=0.05, p=0.96, Cohen’s d=0.02, BF_01_=2.93. There was no statistically significant difference in Glx:GABA+ in the OCC comparison voxel: t(44)=0.62, p=0.54, Cohen’s d=0.19, BF_01_ = 2.89 (See Figure 2C).

## 4 Discussion

Using ^1^H-MRS we tested the hypothesis that there would be PMC GABA and/or Glx and consequent Glx/GABA ratio differences between young APOE E4 carriers and non-carriers measured in the resting state. Specifically, we predicted APOE E4 carriers would show lower PMC GABA+ and/or higher Glx and Glx/GABA+ ratio than non-carriers, resulting in altered E/I balance favouring excitation. This was based on converging lines of evidence: previous fMRI studies of young APOE E4 carriers showing PMC hyperactivation/reduced deactivation (Filippini et al., 2009; Persson et al., 2008; Shine et al., 2015): fMRI-^1^H-MRS findings that levels of PMC deactivation are related to local Glx, GABA+ and resultant E/I balance (Gu et al., 2019; Hu et al., 2013); and circuit-based models of AD, based on studies of transgenic AD mice, which suggest that early hyperactivity is linked to alterations in local neurotransmitters, causing physiological E/I imbalance and network hyperexcitability (Andrews-Zwilling et al., 2010; Busche & Konnerth, 2016; Li et al., 2009; Nuriel et al., 2017; Palop & Mucke, 2016).

Counter to our predictions, our results provide evidence to support the null hypothesis of no difference in PMC (or OCC) neurotransmitter levels between APOE E4 carriers and non-carriers. This was found using frequentist statistics (p > 0.7 in PMC), supported by very low effect sizes (Cohen’s d<0.1 in PMC) and importantly by Bayes factors in favour of the null of ~ 3 (BF_01_ range from 2.9 to 3.2 in PMC). The Bayes factor (BF_01_) provides a continuous measure of evidence for H0 over H1 (and vice versa) (Dienes, 2014; Dienes & Mclatchie, 2018). While there are no necessary thresholds (in contrast to the fixed significance levels of the frequentist approach) to interpret BFs, several authors have suggested that a BF_01_ > 3 is “substantial” evidence in favour of the null (Dienes & Mclatchie, 2018). Therefore, despite the relatively modest sample size, our findings have considerable evidential value (Dienes, 2014; Dienes & Mclatchie, 2018).

There are several potential explanations for our null results, some of which may reflect limitations of ^1^H-MRS studies in general, while some of which may have important implications for the spatial and temporal evolution of AD-related biomarkers across the lifespan in APOE E4 carriers (Harris et al., 2020; Jack et al., 2013; Jagust & Mormino, 2011).

Turning first to the implications of our study for APOE and AD, our finding of no difference between PMC Glx, GABA+ and E/I balance in APOE groups could inform the timeline of spatiotemporal evolution of hyperactivity differences in APOE E4 carriers. We predicted that alterations in E/I balance would account for the previously established alterations in PMC deactivation seen in young adult E4 carriers (e.g. Shine et al., 2015), based on transgenic APOE mouse work showing, for example, that APOE E4 contributes to neuronal hyperactivity by diminishing inhibitory tone, even independently of amyloid (and tau) pathology (Bi et al., 2020; Hijazi et al., 2020; Jiménez-Balado & Eich, 2021; Nuriel et al., 2017). However, detecting no difference in PMC neurotransmitters in young APOE E4 carriers indicates that task-related PM network hyperactivity may not straightforwardly reflect a difference in underlying Glx and/or GABA+ levels. For example, Najm et al. (2019) propose that tau pathology (which may be present even in young adulthood - Braak & Del Tredici, 2011) and mitochondrial impairment occur prior to GABA-ergic cell loss, which then results in E/I imbalance and network hyperexcitability. Other work suggests a complex and potentially synergistic relationship between accumulation of soluble amyloid oligomers, tau and hyperactivity (Busche & Konnerth, 2015; Harris et al., 2020; Palop & Mucke, 2016; Zott et al., 2018). Thus perhaps early subtle AD-related pathology leads to reduced PMC deactivation via fMRI and contributes to a shift in E/I balance, which further contributes to a complex bidirectional relationship between hyperactivity and pathology that only subsequently is detectable via ^1^H MRS later in the lifespan (Quevenco et al., 2019; Schreiner et al., 2016).

Relatedly, APOE E4 may also impact PMC function independent of local neurotransmitter levels and E/I balance. Such factors may include cerebrovascular dynamics (e.g. cerebral blood flow and volume (CBF and CBV), cerebrovascular reactivity (CVR), energy consumption by neurons and glia, and neuronal firing dynamics (rate, amplitude, frequency, phase) (Ekstrom, 2021; Logothetis, 2002; Singh, 2012; Wise et al., 2013)). PMC cerebrovascular reactivity could be a good avenue for further investigation, given alterations in older APOE E4 carriers and increased permeability of the blood-brain-barrier in AD and APOE E4 carriers (Korte et al., 2020; Montagne et al., 2020; Tai et al., 2016; Thambisetty et al., 2010). Two previous studies on the vasculature have detected that young APOE E4 carriers have lower whole brain grey matter CBF (Chandler et al., 2019), and lower hippocampal CO2-CVR in a memory-encoding task during a CO2 challenge (Suri et al., 2015), but the potential impact of such changes on PMC deactivation has yet to be investigated.

Perhaps the most likely explanation for our null finding is that the PMC hyperactivation (reduced deactivation) seen in young APOE E4 carriers reflects not local cortical E/I imbalance, but a *downstream* consequence of initial hyperactivity elsewhere, specifically the medial temporal lobe (MTL). There is dense reciprocal structural and functional connectivity between the MTL and PMC (Bubb et al., 2017; Parvizi et al., 2006; Wang et al., 2016) and network interactions between these regions are important for episodic memory (Ritchey & Cooper, 2020). Network-based accounts of AD suggest that hippocampal hyperexcitability can impact connected PMC regions (e.g. Pasquini et al., 2019) and that AD pathology may spread from the MTL to posterior DN regions via PM network connectivity (Braak & Braak, 1991, 1995; Seeley, 2017). Given evidence of hippocampus and entorhinal cortex hyperactivity on fMRI in young APOE E4 carriers (Filippini et al., 2009; Kunz et al., 2015), as well as increased parieto-temporal connectivity on magnetoencephalography (MEG) (Koelewijn et al., 2019), alongside evidence that APOE E4 mice show prominent early entorhinal cortex hyperactivity linked to reduced inhibitory tone (Nuriel et al., 2017), it would be interesting to study the effect of hippocampal E/I balance on both hippocampal BOLD and its distal effect on PMC BOLD in young APOE E4 carriers. Some evidence suggests that resting hippocampal Glx levels can influence patterns of cortico-hippocampal connectivity on fMRI (Nikolova et al., 2017; Wagner et al., 2016) but the influence of APOE E4 is not yet known. The hippocampus is, however, a challenging brain region to study via ^1^H-MRS, as it suffers from lower SNR due to smaller voxel size required for spatial specificity, and large susceptibility effects leading to broader linewidths and lower spectral resolution (Bednařík et al., 2015). Future ^1^H-MRS studies in the hippocampus could benefit from higher magnetic field strength (e.g. 7T) to improve SNR and spectral quality.

Turning next to ^1^H-MRS limitations, our null finding could in part be due to ^1^H-MRS at 3T not being sufficiently sensitive or specific enough to detect true PMC GABA+ or Glx difference between APOE groups. ^1^H-MRS quantified Glx and GABA+ indicate the metabolite concentrations within the MRS voxel, whereas other aspects of neurotransmission, including glutamate or GABA receptor density at the synapse or binding to receptors, are not accounted for. Such processes cannot be measured using MRI, but could be examined using molecular imaging with PET (e.g. Cuypers et al., 2021). In addition, the Glx and GABA+ measured via ^1^H-MRS indicates MRS-visible pools, which may not all be involved in neurotransmission, as these may include Glx and GABA pools that have a role in metabolism (Kauppinen et al., 1994; Rae, 2014; Stagg & Rothman, 2014). It has therefore been suggested that MRS-measured Glx and GABA+ should be interpreted as excitatory and inhibitory tone, rather than an exact measure of neuronal activity at the time of scanning (Farrant & Nusser, 2005; Harris et al., 2015; Rae, 2014). Having said this, tonic (in contrast to phasic) excitation and inhibition represented by ^1^H-MRS Glx and GABA+ are in fact highly relevant in our study, as they represent activity in the resting state and young APOE E4 carriers show PMC hyperactivity in resting state fMRI scans (Busche & Konnerth, 2016).

In addition, the spatial specificity of MRS (e.g. 2×2×3cm voxel) is much lower than fMRI, where a voxel is measured in millimetres. Any subtle changes in neurotransmitter levels arising in young E4 carriers may be diluted over the large voxel area. Indeed, there is evidence for the co-existence of functional unity but also diversity within the PMC (Parvizi et al., 2006; Yang et al., 2014). Large MRS voxel sizes are required to achieve a sufficient signal-to-noise for metabolites present at low concentrations. This is particularly challenging for GABA+, as its concentration referenced to water tends to be below 3mM (Mullins et al., 2014; Rae, 2014). That said, ^1^H-MRS studies do show functionally relevant correlations between PMC Glx, GABA+ and E/I balance and DN deactivation (Gu et al., 2019; Hu et al., 2013), indicating ^1^H-MRS is sensitive to quantify the metabolites that relate to the BOLD signal. Moving to stronger magnetic fields, such as 7T, could improve sensitivity and reduce voxel size, through improving SNR and resolution of metabolites with overlapping signals (Pradhan et al., 2015; Terpstra et al., 2016).

### Limitations

Unfortunately, several spectra were rejected due to not meeting our data quality criteria. This was particularly the case for PMC MEGA-PRESS spectra, which affected the sample sizes for GABA+ and Glx:GABA+. This was likely due to the smaller PMC voxel size, as fewer GABA spectra were rejected in the larger OCC voxel. The reduced voxel size was important to improve spatial specificity however, so increasing the PCC voxel size would not be a good way to address this. As discussed above, moving to 7T may reduce data exclusion as there is improved SNR at higher magnetic field strength. In future studies, when deciding on sample size we would recommend allowing for the exclusion of more MEGA-PRESS than PRESS spectra.

A reduction in sample size has an impact on power to detect our expected effects. To address this, a sensitivity analysis performed in G*Power reveals that updating the PMC sample sizes to 19 E4 carriers vs 26 non-carriers for Glx, 14 vs 18 for GABA+ and 14 vs 17 for Glx/GABA+ would have 80% power to detect an effect at p<0.05 given an effect size of Cohen’s d = 0.86, 1.03 or 1.05 respectively. These effect sizes are within the effect size of the previously detected PMC deactivation difference between APOE groups in a comparable population (Shine et al., 2015). Thus, despite reductions in sample size, we retained adequate power to detect a large effect between groups. Furthermore, our use of Bayes factors allowed us to assess the extent that our data provide evidence for or against the null hypothesis. Although replicable and precise results are more likely when statistical power is high (Button et al., 2013), it is entirely possible for even low-power experiments to have high evidential value, and contrastingly, for high-power experiments to have low evidential value (Dienes & Mclatchie, 2018). As discussed earlier, Bayes factors in the PMC voxel of interest showed that the data provide substantial evidence in favour of the null (Dienes, 2014).

An improvement to our study that would further inform whether the PM network hyperactivation seen young APOE E4 carriers is related to shifted E/I balance would be to assess both fMRI and/or MEG and ^1^H-MRS within the same participants. Although reduced DN deactivation is a consistent fMRI signature in APOE E4 carriers, correlating BOLD and metabolites in the same participants would provide more direct evidence of whether/how these factors are related. Moreover, a recent development in ^1^H-MRS literature which may reveal more subtle relationships between fMRI and ^1^H-MRS metabolites is functional ^1^H-MRS (fMRS), which here may detect task-related metabolite differences between APOE E4 carriers and non-carriers that are not possible to assess using conventional ^1^H-MRS collected at rest (Apšvalka et al., 2015; Huang et al., 2015; Mullins, 2018; Thielen et al., 2018). In addition, extending the age range of participants, or using a longitudinal design, would further enable us to track how DN deactivation, PMC metabolites and episodic cognition may change with age in APOE E4 carriers to give greater insight into the timeline of how APOE E4 possession may predispose to development of AD.

### Summary

This study provides evidence against differences in PMC Glx, GABA+ and E/I balance in young adult APOE E4 carriers and non-carriers. This suggests that the hyperactivation (or reduced deactivation) consistently observed in this region in APOE E4 carriers is unlikely to be directly associated with altered levels of local neurotransmitters, despite there being evidence in fMRI-^1^H-MRS studies that local Glx, GABA+ and E/I balance are related to PMC deactivation (Gu et al., 2019; Hu et al., 2013). Our null findings could inform models of the spatio-temporal order of alterations within the PMC of APOE E4 carriers that predispose to earlier amyloid accumulation in this region and ultimately development of AD, suggesting that resting PMC neurotransmitter differences do not occur as early as PMC functional changes that are observed in young adult APOE E4 carriers in this region. Alternatively, instead of local effects, it could suggest that PMC hyperactivation in APOE E4 carriers may be related to altered E/I balance elsewhere in the PM network, a key candidate region being the MTL. Our study thus identifies areas for future investigation to gain better understanding of how and why activity differences in PMC occur in young APOE E4 carriers that subsequently predispose to earlier AD pathology.

## Supplementary Material

### 5 MRS pilot study: Test-retest reliability

#### 5.1 Introduction

The purpose of the pilot study was to assess the test-retest reliability of the six main ^1^H-MRS metabolites. In the PRESS scan these were Glx, N-acetyl-aspartic-acid (tNAA), myo-Inositol (mI), creatine (tCr) and choline (Cho), and GABA+ in the MEGA-PRESS scan.

#### 5.2 Methods

Six participants (3 male, 3 female, mean age 25.5 ± 1.6 years) took part in two scan sessions. The study was approved by the Cardiff University School of Psychology research ethics committee, and written informed consent was obtained from all participants. As in the main text, scans were scheduled for the female participants in the luteal phase of the menstrual cycle, to avoid any effects of menstrual cycle phase on metabolite values. Scans were scheduled an average of 3 weeks apart.

Scans were performed at the Cardiff University Brain Research Imaging Centre (CUBRIC) on a 3T General Electric (GE) HDx scanner fitted with an 8-channel phased array head coil. The two scan sessions for each participant included a structural scan, two PCC PRESS scans, two OCC PRESS scans, and two PCC MEGA-PRESS scans. This meant that each scan type was repeated within a scan session, allowing for an intra-scan comparison of metabolite levels when the voxel was placed in the same position, and an inter-scan comparison between scan sessions. As in the main text, the PCC voxel measured 2×2×3cm, and the OCC voxel 3×3×3cm (see Figure 1). The same PRESS scan parameters as in section 2.4 were used, and the PCC MEGA-PRESS scans used the parameters TE = 68ms, TR = 1800ms, number of averages = 332, spectral width 5kHz, number of data points = 4096, acquisition time = 10minutes (i.e. a shorter scan time with fewer averages than in the main text). We did not measure GABA+ in OCC MEGA-PRESS scans, as a test-retest reliability assessment had been done previously in CUBRIC, and a 3×3×3cm OCC voxel was a standard method at this centre (NB. Our PCC MEGA-PRESS scan parameters matched that of the OCC voxel in these studies) (Mikkelsen et al., 2015; Muthukumaraswamy et al., 2009, 2012).

Data analysis was the same as described in Section 2.5. Briefly, PRESS data (for Glx, tNAA, mI, TCr, Cho) were analysed using TARQUIN (Totally Automatic Robust Quantification In NMR) version 4.3.3 (Wilson et al., 2011). MEGA-PRESS data (for GABA+) were analysed using GANNET (GABA-MRS Analysis Tool) version 2.0 (Edden et al., 2014). Metabolite signals were referenced to H_2_O, corrected for the fraction of CSF in the voxel, and expressed as a concentration in millimoles (mM) per unit tissue volume. Data were excluded where the CRLB was above 20.

To test the intra-scan test-retest reliability, the coefficient of variation (CV) was calculated for the pairs of scans within each scan session (i.e. for scan 1a and 1b; and for scan 2a and 2b). For the inter-scan test-retest reliability, the coefficient of variation was calculated for the first scan in each session (i.e. for scan 1a and 2a). CV is calculated as (standard deviation/average x 100). Metabolite values from the four scans were also compared to assess if they were significantly different from each other, using a repeated measures ANOVA (SPSS).

#### 5.3 Results and Discussion

The following data were excluded due to CRLB>20: in the OCC voxel, two Cho, two Glx, three mI, two tCr and two tNAA values; in the PCC voxel, one mI value.

Supplementary Table 1 lists the intra- and inter- scan CVs. There were no significant differences in any metabolite across the four scans (see Supplementary Table 1; all p>0.08).

Previous studies of PRESS scan test-retest reliability have reported CVs of 5% for tNAA, Cr and mI, and 9% for tCho in the cortex measured at 1.5T (Geurts et al., 2004), or 7% for tNAA, Cr and tCho, and 10% for mI at 3T (Kirov et al., 2012), and have suggested these values represent a reliable measurement. The CV for GABA+ quantified from a MEGA-PRESS scan acquired at 3T was 4% in the OCC and 14.8% in an anterior cingulate cortex voxel (OCC voxel measured 3×3×3cm, ACC voxel 2×3×4cm) (Mikkelsen et al., 2015). Our CVs are comparable to these findings.

The ability of Gannet to fit the GABA+ data to the model was noted to be lower for the PCC GABA+ data than found for OCC GABA+ data in CUBRIC, indicated by larger residuals in the PCC data. Therefore, to improve PCC GABA+ data quality in our main study, the scan time was lengthened from 10 to 15 minutes to acquire an extra 180 averages.

**Supplementary Table 1:**
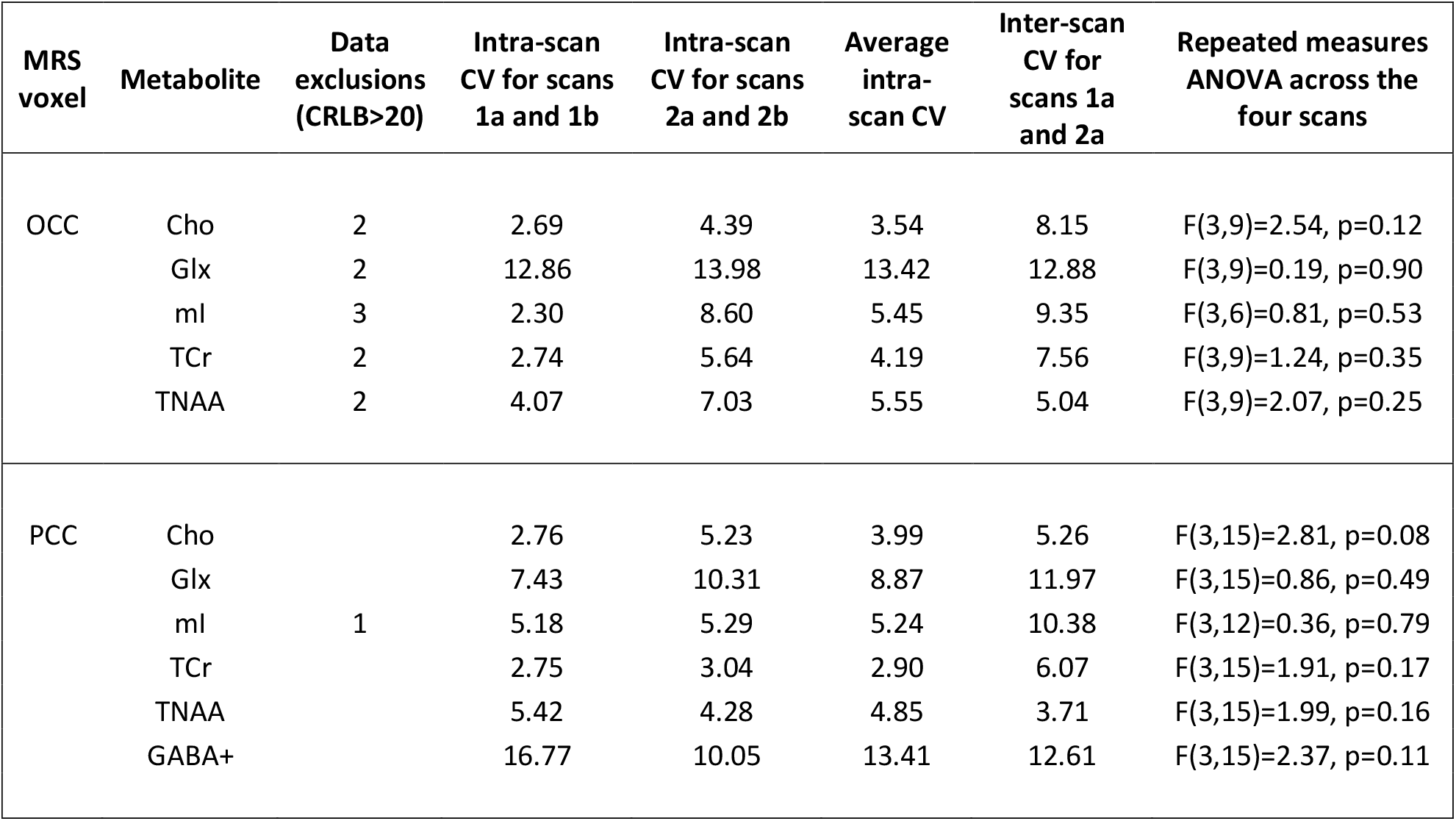
Intra- and inter-scan coefficient of variation (CV) and statistical analysis across four scans to assess test-retest reliability of PRESS metabolites in the OCC and PCC voxels, and of MEGA-PRESS GABA+ in the PCC voxel.

## 6 Comparison of other main ^1^H-MRS metabolites: replication of Suri et al. (2017)

### 6.1 Introduction

The ^1^H-MRS scan sequence with a TE of 35ms used to quantify Glx in the main text also allows for quantification of a range of metabolites commonly studied in ^1^H-MRS papers. The main metabolites interest, which are the most often studied and of greatest relevance to APOE E4 carriers as they are altered in AD, are N-acetyl-aspartic-acid (tNAA), myo-Inositol (mI), creatine (tCr) and choline (Cho).

NAA is interpreted as a marker of neuronal density and integrity, as it is synthesized predominantly in neuronal mitochondria (Patel & Clark, 1979; Rae, 2014). It is considered a neuronal marker as tNAA is negatively correlated with atrophy in neurodegenerative disorders (Oz et al., 2010, 2014), and positively correlated with synapse density in a ^1^H-MRS study of elderly participants followed by histology on post-mortem tissue (Murray et al., 2014). The “t” in tNAA denotes “total”, as the large NAA peak in the MRS spectrum has a small peak on its shoulder from N-acetyl-aspartic glutamate (NAAG), which has a highly similar chemical structure, so is difficult to accurately separate from NAA (Edden et al., 2007). Therefore, these two signals are combined and reported together as standard.

MI is considered to be a marker of inflammation. It is associated with gliosis, the recruitment and proliferation of glial cells to an affected region as a response to injury or disease (Rae, 2014). This was established first in cancer (glioma) and subsequently in other neuroinflammatory disorders such as multiple sclerosis, as well as acute inflammation due to traumatic brain injury (Oz et al., 2014).

Similar to TNAA, tCr is the sum of the peaks on the MRS spectrum from Cr and phosphocreatine (PCr), which is the product of the phosphorylation of Cr by creatine kinase (Rae, 2014). This reaction is termed the creatine kinase/phosphocreatine (CK/PCr) energy shuttle, and is an important way that the cell replenishes its supply of ATP. TCr, therefore, is used as a marker of energy metabolism.

The Cho signal represents several choline containing compounds, which include free choline (Cho), phosphocholine (PC) and glycerophosphocholine (GPC) (Barker et al., 1994; Miller et al., 1996). Membrane-bound choline in phospholipids is MRS-invisible (Miller et al., 1991), thus the signal obtained in MRS represents cytosolic choline compounds (Stagg & Rothman, 2014). In MRS, the Cho peak is considered a marker of cell membrane integrity or turnover (Rae, 2014).

In ^1^H-MRS studies of AD patients, a decrease in PMC tNAA and an increase in PMC mI is a robust finding replicated several times (Kantarci et al., 2000, 2007, 2013; Murray et al., 2014; Voevodskaya et al., 2016; Wang et al., 2015). The increase in PCC mI may precede the decrease in tNAA, as one study found that increased mI was detected in both AD and MCI patients, while decreased PCC tNAA was only detected in AD patients (Kantarci et al., 2000). This suggests AD is characterized by an increase in inflammation and a decrease in neuronal density in this brain region. Changes in Cho are less consistent, but there are reports of an elevation in Cho (Kantarci et al., 2004, 2007). Many studies use Cr as a reference for other metabolites, so tCr tends not to be reported alone, however this is cautioned against as any tCr changes in AD could lead to misleading results for the other metabolites (Rae, 2014; Stagg & Rothman, 2014).

Some studies have detected ^1^H-MRS metabolite changes in older APOE E4 carriers in the PMC. Differences include higher mI in carriers compared to non-carriers (Voevodskaya & Sundgren, 2016), higher mI/Cr and Cho/Cr (Gomar et al., 2014), lower Cr (Laakso et al., 2003) and lower tNAA (Riese et al., 2015). This pattern of metabolite differences between older E4 carriers and non-carriers have not always been detected however (Kantarci et al., 2002; Laakso et al., 2003; Suri et al., 2017).

A key study of note here is by Suri et al. (2017) which analysed PMC metabolites in a group of old (age 60-85) and young participants (age 20-40), which was also split into APOE E4 carriers and non-carriers. They studied a 2×2×2cm PMC voxel, placed in a highly similar location to our PMC voxel. They found a significant effect of age on certain metabolites, where older participants had higher PCC mI and tCr, and lower PMC glutamate. However, there was no significant effect of APOE group or interaction of APOE group and age on any metabolite. Although this paper had a large sample size in their older group (n=117 participants, inc. 22 E4 carriers), a limitation of the study is that their younger group only had a sample size of 8 APOE E4 carriers compared to 22 non-carriers. This could limit the power to detect an effect, as it is lower than the sample size often used in MRS studies people at genetic risk of disease with controls (e.g. 22 participants at genetic risk of schizophrenia vs 22 controls (Yoo et al., 2009) and 23 risk participants vs 24 controls (Tandon et al., 2013)).

The purpose of our analysis was to use existing data from our Glx analysis to replicate Suri et al. (2017), testing whether young APOE E4 carriers have any differences in the four main metabolites studied in AD compared to non-carriers (tNAA, mI, tCr, Cho). Advantages of our study are that we had a larger sample size of young E4 carriers (n=20 carriers vs 27 non-carriers), and we included an OCC voxel as a control to test whether any metabolite differences were region specific.

### 6.2 Methods

The methods are the same as in the main text. The four metabolites analyzed here were quantified in the same analysis of the PRESS scan data for Glx using Tarquin v4.3.3, but not included in the main text as our focus there was on Glx and GABA.

Data excluded due to CRLBs over 20 were, in the PMC, two tNAA (two E4 carriers), 14 mI (four E4 carriers, 10 non-carriers), two tCr (one E4 carrier, one non-carrier), three Cho (one E4 carrier, two non-carriers), and in the OCC, one tNAA, four mI, one tCr and two Cho (all E4 carriers).

### 6.3 Results and Discussion

There were no differences in the concentrations of the four metabolites between APOE E4 carriers in and non-carriers in the PMC voxel of interest: tNAA, t(42)=0.85, p=0.40, Cohen’s d = −0.26, BF_01_=2.49; mI, t(31)=0.70, p=0.49, Cohen’s d= −0.24, BF_01_=2.49; tCr, t(43)=0.03, p=0.97, Cohen’s d= −0.01, BF_01_=3.36; tCho, t(42)=0.62, p=0.54, Cohen’s d= −0.19, BF_01_=2.86.

There were also no differences between E4 carriers and non-carriers in the OCC comparison voxel: tNAA, t(44)=1.09, p=0.28, Cohen’s d = −0.33, BF_01_=2.09; mI, t(41)=0.0005, p=1.00, Cohen’s d= 0.0001, BF_01_=3.24; tCr, t(44)=0.93, p=0.36, Cohen’s d= −0.28, BF_01_=2.39; tCho, t(43)=0.97, p=0.34, Cohen’s d= −0.29, BF_01_=2.30.

Our PCC findings therefore replicate the null findings of Suri et al. (2017) in the younger APOE E4 carriers and suggest that findings in older adults may reflect consequences of aging/disease related processes.

---

**Supplementary Figure 1:**
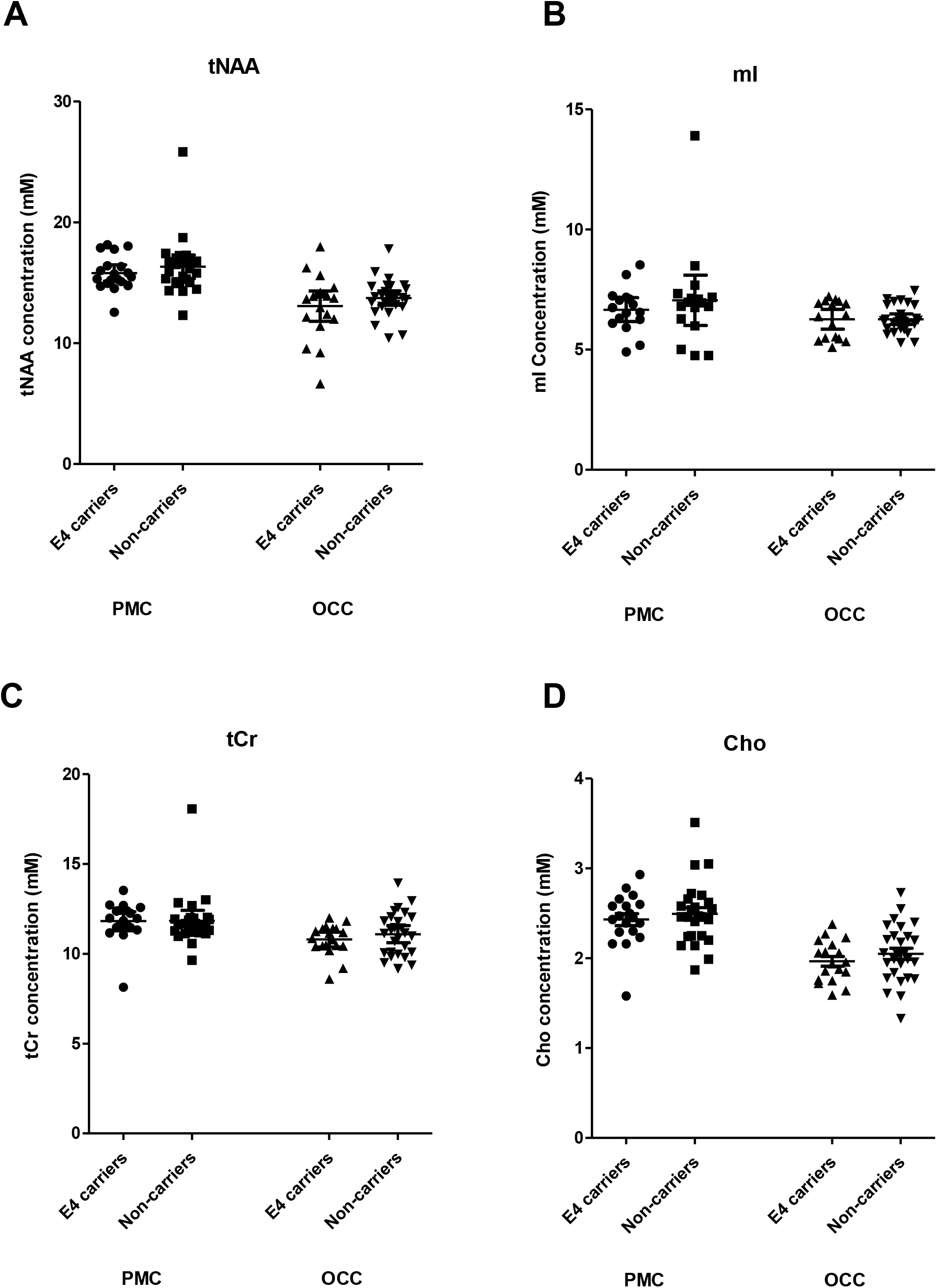
Comparison of ^1^H-MRS metabolites in APOE E4 carriers and non-carriers in the PMC voxel of interest and OCC comparison voxel. Plots show individual data points, lines represent the mean, and error bars are 95% confidence intervals.

## Acknowledgements

We are grateful to Mark Mikkelsen for sharing Matlab scripts for ^1^H-MRS analysis, and to Peter Hobden for assistance with scanning. The work was funded by the Wellcome Trust (Strategic Award – KSG, ADL, 104943/Z/14/Z) and Medical Research Council (KSG, G1002149). AGC’s PhD studentship was funded by the Cardiff University Neuroscience and Mental Health Research Institute.

## CRediT author statement

**Costigan AG:** Conceptualisation, Investigation, Formal analysis, Visualization, Writing-Original Draft; **Umla-Runge, K:** Conceptualisation, Investigation, Project administration; **Evans, CJ:** Methodology, Software, Validation; **Raybould R:** Investigation; **Graham, KS:** Conceptualization, Project administration, Supervision; **Lawrence, AD:** Conceptualisation, Supervision, Writing – Review and Editing

